# Dephosphorylation is the Mechanism of Fibroblast Growth Factor Inhibition of Guanylyl Cyclase-B

**DOI:** 10.1101/191684

**Authors:** Jerid W. Robinson, Jeremy R. Egbert, Julia Davydova, Hannes Schmidt, Laurinda A. Jaffe, Lincoln R. Potter

## Abstract

Activating mutations in fibroblast growth factor receptor 3 (FGFR3) and inactivating mutations of guanylyl cyclase-B (GC-B, also called NPRB or NPR2) cause dwarfism. FGF exposure inhibits GC-B activity in a chondrocyte cell line, but the mechanism of the inactivation is not known. Here, we report that FGF exposure causes dephosphorylation of GC-B in rat chondrosarcoma cells, which correlates with a rapid, potent and reversible inhibition of C-type natriuretic peptide-dependent activation of GC-B. Cells expressing a phosphomimetic mutant of GC-B that cannot be inactivated by dephosphorylation because it contains glutamate substitutions for all known phosphorylation sites showed no decrease in GC-B activity in response to FGF. We conclude that FGF rapidly inactivates GC-B by a reversible dephosphorylation mechanism, which may contribute to the signaling network by which activated FGFR3 causes dwarfism.

**Highlights:**

1. Guanylyl Cyclase-B is expressed in rat chondrosarcoma cells
2. FGF2 induces a rapid, potent, and reversible inhibition of GC-B
3. FGF2 treatment causes GC-B dephosphorylation
4. FGF2 does not inhibit a dephosphorylation-resistant form of GC-B
5. Dephosphorylation is the mechanism of FGF2-dependent inhibition of GC-B

**Abbreviations:**

cGMP: cyclic guanosine monophosphate
GC: guanylyl cyclase
NP: natriuretic peptide
PBS: phosphate buffered saline
WT: wild type

## 1. Introduction

C-type natriuretic peptide (CNP) is a paracrine factor that stimulates the growth of long bones and vertebrae, promotes axonal bifurcation in the spinal cord, and prevents resumption of meiosis in the ovarian follicle [1-3]. These physiologic functions of CNP are mediated by guanylyl cyclase-B (GC-B), which elevates intracellular cGMP in response to CNP binding. Female mice lacking GC-B are infertile, and mice of both sexes lacking functional CNP or functional GC-B exhibit disproportionate dwarfism caused by reduced chondrocyte proliferation and hypertrophy [4, 5]. In humans, genetic mutations that inactivate both alleles encoding GC-B cause acromesomelic dysplasia, type Maroteaux (AMDM) dwarfism [6]. Conversely, mutations that increase CNP expression [7, 8] or mutations that activate a single GC-B allele in the absence of CNP cause skeletal overgrowth [9-11]. CNP levels in plasma are also predictive of longitudinal bone growth [12, 13].

GC-B is a single membrane-spanning enzyme that exists as a higher ordered oligomer, possibly a dimer, that catalyzes the synthesis of cGMP from GTP in response to CNP binding [14]. The extracellular domain of GC-B is glycosylated and terminal N-linked glycosylation is required for the formation of an active GC catalytic domain [15, 16]. AMDM dwarfism-causing missense mutations are most often associated with receptors lacking terminal N-linked glycosylation that markedly reduces or abolishes the ability of GC-B to form an active catalytic domain [16]. The intracellular portion of GC-B consists of a kinase homology domain that contains six chemically identified serine/threonine phosphorylation sites and one putative, functionally identified serine phosphorylation site [17-19], a short coiled-coiled dimerization region, and a carboxyl-terminal GC domain [14, 20].

Previous studies have established that phosphorylation of GC-B is required for activation of GC-B. Conversion of all six chemically determined serine and threonine phosphorylation sites in GC-B to alanines produced a properly folded enzyme that retained GC activity under synthetic detergent activation conditions but had only 6% of the CNP-dependent activity observed with the phosphorylated wild type enzyme [17]. Conversely, mutating all chemically identified phosphorylation sites and one putative functionally identified phosphorylation site to glutamate to mimic the negative charge of phosphate produced an enzyme called GC-B-7E that is activated by CNP like the phosphorylated WT enzyme [19].

Early studies showed that activation of several G protein coupled receptors inactivates GC-B, through a process involving dephosphorylation [21, 22]. Recently, luteinizing hormone (LH) was shown to stimulate GC-B dephosphorylation and inactivation in ovarian follicles by a process that requires a PPP family serine and threonine protein phosphatase [23]. Importantly, a knock-in mouse (GC-B^7E/7E^) expressing GC-B-7E at the normal GC-B genetic locus was immune to LH-dependent GC-B inactivation and displayed a 5-h delay in the resumption of meiosis in the oocyte [24]. These findings indicate that hormones that activate G protein coupled receptors inactivate GC-B by dephosphorylation.

The present paper investigates the possibility that not only GPCR signaling, but also growth factor receptor signaling could inactivate GC-B by dephosphorylation. Multiple mechanisms could contribute to FGFR3 regulation of long bone growth [25], including FGF inhibition of GC-B [26]. However, although FGF2 exposure was shown to inactivate GC-B in the ATDC5 chondrocyte cell line [26] and in BALB/3T3 fibroblasts [27], the molecular basis for the inactivation was not determined. Here, we used multiple approaches to examine the molecular mechanism of FGF2-dependent GC-B inactivation in rat chondrosarcoma (RCS) cells, a highly physiologic chondrocyte cell line [28].

## 2. Material and methods

### 2.1. Materials

^125^I cGMP radioimmunoassay kits were from PerkinElmer Life Sciences (Waltham, MA). CNP and heparin were from Sigma-Aldrich (St. Louis, MO), and FGF2 was from R&D Systems (Minneapolis, MN). IPA300 Protein A-conjugated resin was from Repligen (Waltham, MA, USA).

### 2.2. Cell culture

RCS cells are derived from a Swarm rat chondrosarcoma and express FGFR2 and FGFR3, but the mRNA for FGFR3 is at least seven-fold higher than the mRNAs of the other FGF receptor [28-33]. The RCS cells were a gift from Professor Benoit de Crombrugghe (MD Anderson Cancer Center, Houston, TX) and were maintained in DMEM with 1% penicillin/streptomycin and 10% fetal bovine serum. Except as indicated, FGF2 was used at a concentration 100 ng/m in the presence of 1 μg/ml heparin. Control cells were treated with heparin alone.

### 2.3. Construction of adenovirus-based vectors

The replication-deficient CMV promoter-driven rat GC-B-expressing vectors (RGD-CMV-GC-B-7E and RGD-CMV-GC-B-WT) were constructed through homologous recombination with the RGD fiber-modified Ad backbone plasmid (RGD-Ad-Easy). All vectors are identical and contain the CMV promoter-driven GC-B transgene cassette inserted in place of the deleted E1 region of a common Ad vector backbone. First, full-length GC-B-WT or GC-B-7E sequences derived from pRK5-GC-B were cloned into pShuttle-CMV plasmid. The resultant plasmids were linearized with Pme I digestion and subsequently co-transformed into *E. coli* BJ5183 with the RGD fiber-modified Ad backbone plasmid (RGD-Ad-Easy). After selection of recombinants, the recombinant DNA was linearized with Pac I digestion and transfected into 911 cells to generate viral vectors. The virus was propagated in 911 cells, dialyzed in phosphate-buffered saline (PBS) with 10% glycerol, and stored at -80°C. Titering was performed with a plaque-forming assay using 911 cells and optical density-based measurement.

### 2.4. Adenoviral Transduction

A 10 cm dish of RCS cells at 50% confluency was transduced with either RGD-CMV-GC-B-7E or RGD-CMV-GC-B-WT using a multiplicity of infection of 100. Cells were incubated overnight, followed by a change in medium. GC activity was assayed in membranes from serum-starved cells harvested two days after viral transduction.

### 2.5. GC Assays

Crude membranes were prepared in phosphatase inhibitor buffer as previously described [34]. Assays were performed at 37°C for the indicated times in a cocktail containing 25 mM HEPES pH 7.4, 50 mM NaCl, 0.1% BSA, 0.5 mM isobutylmethylxanthine, 1 mM EDTA, 5 mM MgCl_2_, 0.5 μM microcystin, and 1X Roche Complete protease inhibitor cocktail. Unless indicated, the mixture also included 1 mM ATP and 1 mM GTP. If not indicated otherwise, CNP concentrations were 1 μM. Assays with 1% Triton X-100 and 5 mM MnCl_2_ substituted for MgCl_2_ were used to determine the total amount of GC-B in the membranes, since phosphorylation does not affect GC activity measured in detergent. Reactions were initiated by adding 80 μl of the mixture to 20 μl of crude membranes containing 5-15 μg of crude membrane protein. Reactions were stopped with 0.4 ml of ice-cold 50 mM sodium acetate buffer containing 5 mM EDTA. Cyclic GMP concentrations were determined by radioimmunoassay as described [35].

### 2.6. Immunoprecipitations and ProQ or SYPRO Ruby Staining

RCS cells were lysed for 30 min at 4°C on a rotator in RIPA buffer containing: 50 mM HEPES pH 7.4, 50 mM NaF, 2 mM EDTA, 0.5% deoxycholate, 0.1% SDS, 1% IGEPAL CA-630, 100 mM NaCl, 10 mM NaH_2_PO_4_, 1X Roche Protease Inhibitor Cocktail, and 0.5 μM microcystin. Cellular extracts were then precleared on a rotator in the same RIPA buffer at 4 °C containing 50 μl IPA300 Protein A-conjugated resin for 30 min. Samples were centrifuged and the supernatant transferred to a fresh tube. 25 μl IPA300 Protein A-conjugated resin, and 2 μl anti-GC-B rabbit polyclonal primary antibody 6327 that was immunized against the last 10 C-terminal amino acids of rat GC-B, were added to the samples and rotated over night at 4 °C. The resin was washed three times in RIPA buffer without NaCl or NaH_2_PO_4_, and then resuspended in protein sample buffer and boiled 5 min.

Immunocomplexes of GC-B were fractionated on an 8% SDS polyacrylamide gel, then the gel was sequentially stained with ProQ Diamond followed by SYPRO Ruby dyes as previously described [21, 36]. Densitometry ratios were calculated by dividing the Pro-Q Diamond signal intensity (Phospho-GC-B) by the SYPRO Ruby signal intensity (Processed GC-B, which means processed in the ER by glycosylation) using the LiCor Image Studio software.

### 2.7. Phos-tag gel electrophoresis

For analysis of phosphorylation by Phos-tag, GC-B was immunoprecipitated as previously described [23]. Briefly, ∼200-500 μg crude membrane protein was diluted to 0.5 or 1 ml in 50 mM Tris-HCl pH 7.5, 50 mM NaF, 10 Mm NaH_2_PO_4_, 2 mM EDTA, 0.5% deoxycholate, 0.1% SDS, 1% NP-40, 100 mM NaCl, 10 mM NaH_2_PO_4_, 1X Roche Protease Inhibitor Cocktail, and 1 μM microcystin. After adding 0.6 or 1 μl anti-GC-B rabbit polyclonal antiserum 6328, made against a C-terminal peptide of GC-B [35], samples were rotated at 4 °C for 1 hour, then added to 25 or 50 μl Protein A/G magnetic beads (ThermoFisher Scientific) and rotated overnight at 4 °C. The beads were washed three times in the same buffer and protein was eluted for 10 min at 70 °C in protein gel sample running buffer with 75 mM dithiothreitol.

Phos-tag gel electrophoresis and western blotting were then performed as described, using a primary antibody made against the extracellular domain of GC-B [37]. For Fig. S1, the 6327 antibody against the C-terminus of GC-B was used. The blots were developed with WesternBright Sirius reagent (Advansta, Menlo Park, CA). For densitometry, the amount of staining for the upper, more phosphorylated region was divided by the amount of staining for the lower, less phosphorylated bands and was presented as the phos-tag intensity ratio. Because the absolute values of the ratios were variable across blots, we normalized the data by dividing all of the ratios on a given blot to the mean ratio of the control lanes on that blot.

### 2.9. Statistical Analysis

Statistics and graphs were generated using GraphPad Prism 7. p values were generated using unpaired, two-tailed student’s t-test. Statistical significance was determined where p ≤ 0.05. The vertical bars represent the standard error of the mean. Where error bars are not visible, they are within the symbol.

## 3. Results

### 3.1. RCS cells express GC-B

Whether RCS cells express GC-B, the highly-related receptor GC-A, or both is not known. The presence of individual natriuretic peptide receptors was investigated by examining the ability of low and high concentrations of ANP or CNP to stimulate cGMP production by GC-A or GC-B, respectively, in RCS cell membrane preparations. One hundred nM CNP increased GC activity 2.1-fold and 1000 nM CNP increased activity 5.7-fold over basal levels (Fig. 1). In contrast, 100 nM and 1000 nM ANP increased cGMP by only 1.47 and 1.75-fold, respectively. These data indicate that GC-B is the dominant natriuretic peptide receptor in RCS cells.

**Figure 1.**
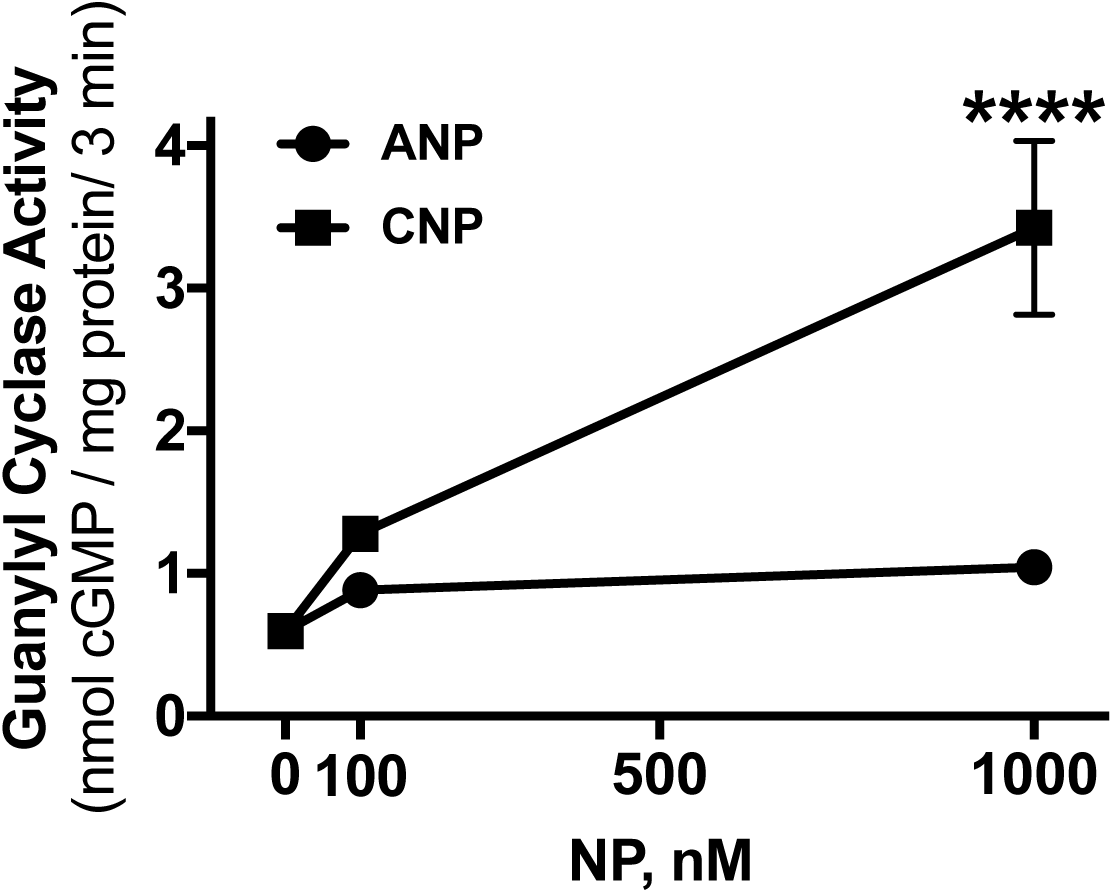
RCS cells express GC-B. Crude membranes from RCS cells were assayed for GC activity with 1 mM ATP in the presence of 100 nM or 1000 nM concentrations of ANP or CNP as indicated. Each point represents the average of duplicate determinations from 3 separate experiments such that n=3 for each point. Symbols represent the mean and the vertical bars represent the standard error. Where error bars are not visible, they are contained within the symbol. **** indicates p<0.0001, using an unpaired, two-tailed student’s t-test.

### 3.2. FGF2 is a potent and rapid inhibitor of GC-B

To determine whether FGF2 inhibits GC-B, RCS cells were incubated for 15 min in the absence or presence of increasing concentrations of FGF2 and 1 μg/ml heparin, its coactivator. Heparin by itself had no effect. Among the many FGF isoforms, FGF2 was used because it activates FGFR3 as well as FGFR2, and older data indicate that both receptors are expressed in RCS cells [32, 38]. However, more recent data by Chapman et al. demonstrated that the mRNA for FGFR3 is 8-fold higher than FGFR1, which is 100-fold higher than FGFR2 and 50-fold higher than FGFR4 [31]. FGF2 inhibited CNP dependent GC-B activity at sub-saturating (100 nM) and saturating (1 μM) CNP concentrations with a similar IC_50_ of approximately 5 ng/ml FGF2 (Fig. 2A). To determine how rapidly FGF2 inhibited GC-B, RCS cells were incubated for increasing periods of time in the presence of 100 ng/ml FGF2, then membranes were prepared and assayed for CNP-dependent GC activity at sub-saturating (100 nM) and saturating (1 μM) CNP concentrations. FGF2 inhibited GC-B activity with a t_1/2_ of ∼7 min, with maximal inhibition occurring by 15 min for both CNP concentrations. The FGF-dependent inhibition of GC-B was sustained for at least 1 h when activated with either concentration of CNP (Fig. 2B).

**Figure 2.**
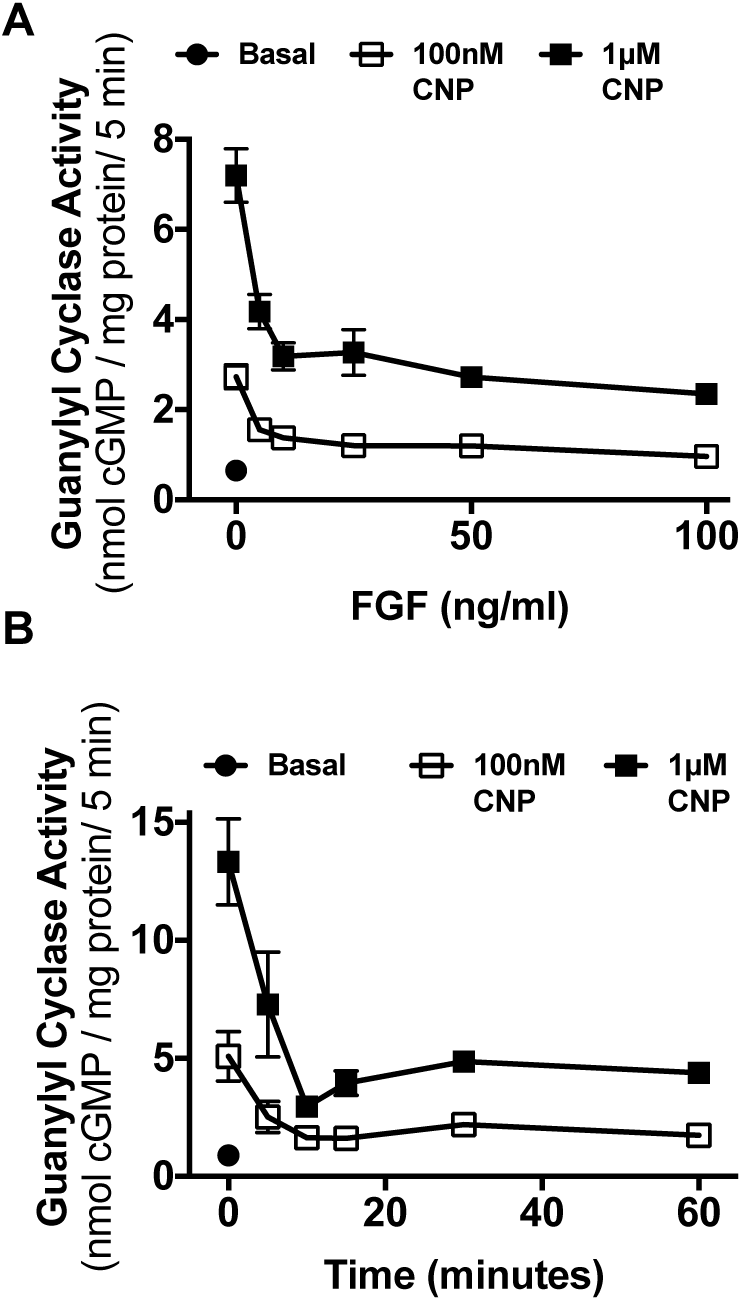
FGF2 is a rapid and potent inhibitor of GC-B in RCS cells. (A) GC activity was measured in the absence of CNP (basal) or in the presence of 100 nM or 1 μM concentrations of CNP in membranes prepared from whole cells that were incubated for 15 min with 1 μg/ml heparin in the presence or absence of the indicated concentrations of FGF2. (B) GC activity was measured in membranes from whole cells treated in the presence or absence of 100 ng/ml FGF2 for the periods of time shown. Each point in (A) and (B) represents the average of duplicate determinations from 3 separate experiments such that n = 3 for each point.

### 3.3. FGF2 inhibition of GC-B is rapidly reversible

The reversibility of the FGF2-dependent inhibition was also investigated. RCS cells were incubated in the presence or absence of 100 ng/ml FGF2 for 15 min followed by two rapid washes with DMEM lacking serum to remove the FGF2. The cells were incubated in the same medium for the indicated periods of time before the medium was aspirated. The cells were immediately removed, sonicated, and crude membranes were prepared and assayed for CNP-dependent GC activity (Fig. 3). FGF2 removal was associated with a rapid re-activation of GC-B with activity being completely recovered by 30 min and remaining elevated above initial levels for at least 2 h.

**Figure 3.**
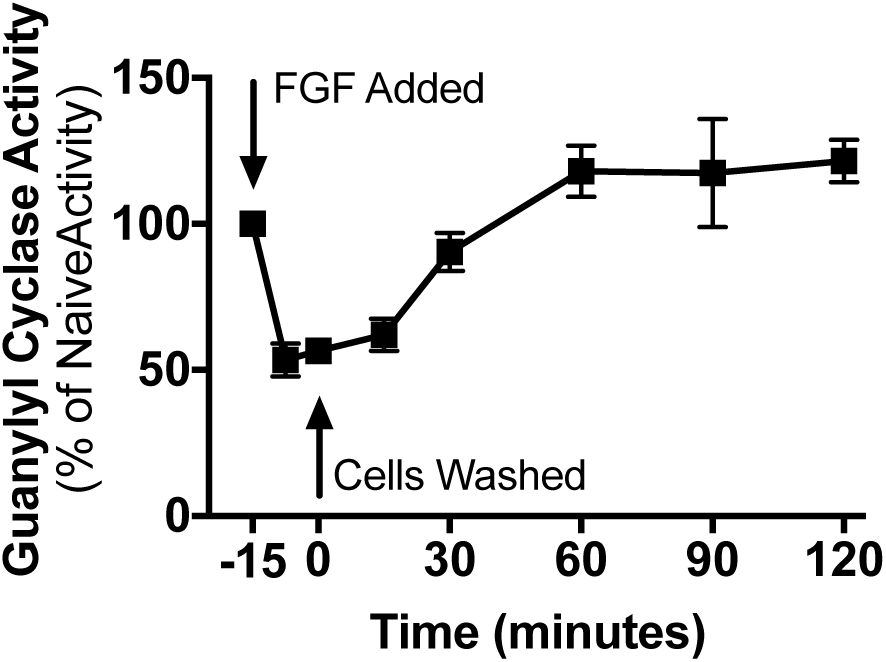
FGF2 inhibition of GC-B is rapidly reversible. GC activity was measured in membranes prepared from whole RCS cells treated with 1 μg/ml heparin and 100 ng/ml FGF2 for 15 min, washed twice, and incubated with fresh cell culture medium at 37°C for the periods of time shown before crude membranes were prepared and assayed for GC activity in the presence of 1 μM CNP. Each point represents the average of duplicate determinations from five separate experiments such that n = 5 for each point, where vertical error bars within the symbols represent the standard error.

### 3.4. FGF2 exposure results in GC-B dephosphorylation

Because inhibition of GC-B by multiple hormones and signaling factors is correlated with receptor dephosphorylation [23, 24, 27, 35, 39, 40], we investigated whether dephosphorylation contributes to the FGF2-dependent inhibition of GC-B in RCS cells. To determine GC-B phosphate levels, we employed two different phosphate detection methods, Phos-tag and ProQ Diamond. Phos-tag phosphate detection uses a dinuclear metal complex to bind phosphoryl groups on proteins to retard their migration through a polyacrylamide gel [23, 41]. The more phosphorylated protein species migrate more slowly and are located higher in the gels, whereas the less phosphorylated species migrate faster and are located lower in the gel. This assay provides a qualitative but highly sensitive indicator of changes in GC-B phosphorylation. In contrast, ProQ Diamond is a small, organic fluorescent dye that non-covalently stains phospho –serine, – threonine, and –tyrosine residues [42]. We previously showed that ProQ Diamond staining is highly correlated with the ^32^PO_4_ content of GC-B isolated from metabolically labeled cells [36]. Thus, the ProQ method is quantitative, but it is less sensitive than the Phos-tag method. SYPRO Ruby staining determined GC-B protein levels on the same gel used for the ProQ Diamond staining to demonstrate that any changes in phosphate were not due changes in protein levels.

The Phos-tag but not the ProQ Diamond method was sufficiently sensitive to detect the phosphorylation of endogenous GC-B in extracts from RCS cells (Fig. 4A and B). In control samples that were not treated with FGF2, GC-B migrated as a broad upper region, and 2 lower bands. After FGF2 exposure, the intensity of the upper region decreased, and the intensity of the lower region increased, indicating dephosphorylation of GC-B. To quantify the change, we measured the ratio of the intensities of the upper and lower regions, normalized to the mean ratio of the control lanes for each blot. FGF2 decreased the ratio of the intensities of the slower migrating, more phosphorylated region versus the faster migrating, less phosphorylated bands by 64% (Fig. 4A,B).

**Figure 4.**
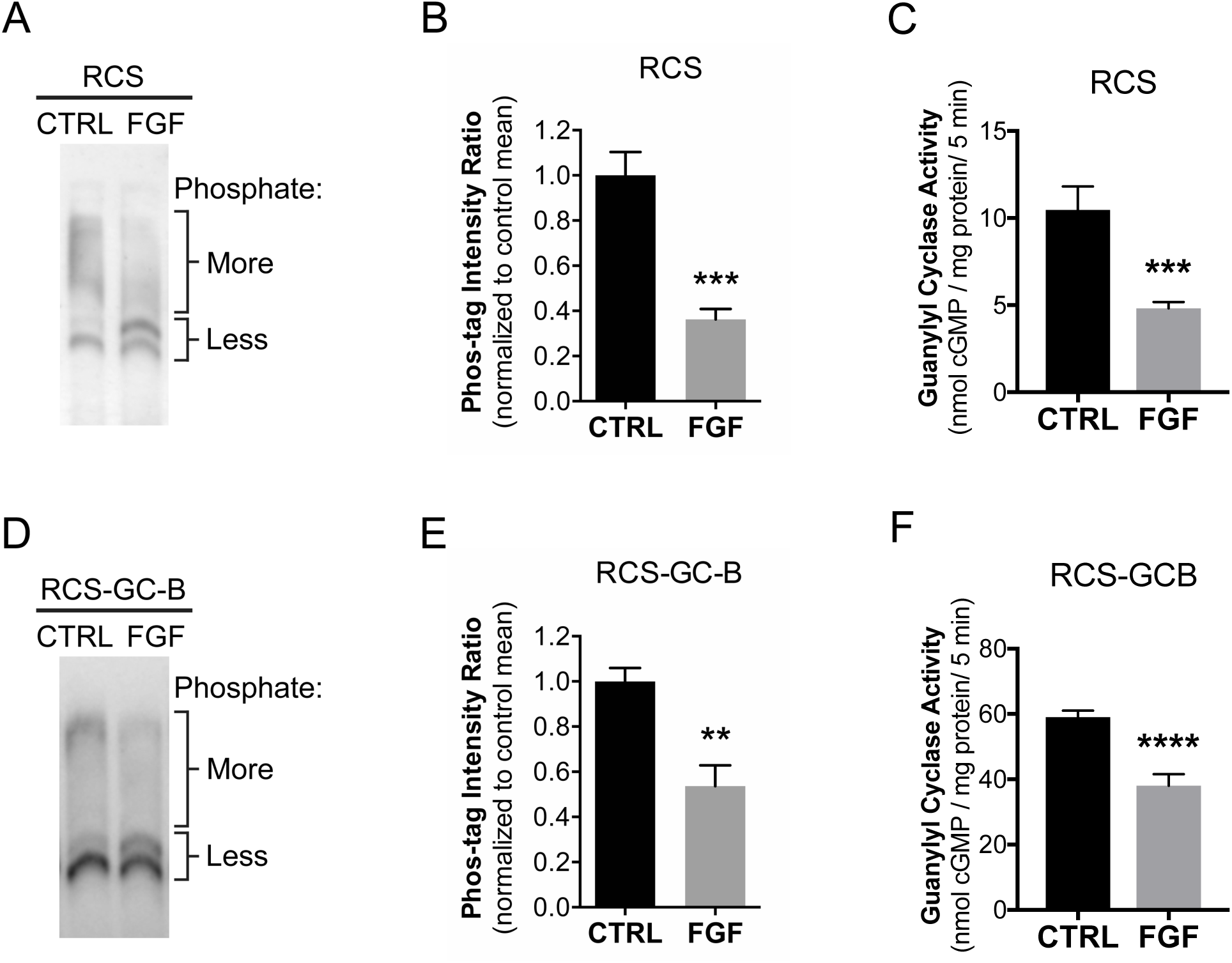
Phos-tag analysis indicates that FGF2 stimulates GC-B dephosphorylation. RCS cells, or RCS cells virally transduced with wild type GC-B (RCS-GC-B) were treated with 1 μg/ml heparin in the absence or presence of 100 ng/ml FGF for 20 min. Cells were lysed and crude membranes were immunoprecipitated with an antibody against the C-terminus of GC-B. The immunocomplexes were washed and fractionated on a Phos-tag gel and western blotted with an antibody against the extracellular domain of GC-B. (A-C) RCS cells that endogenously express GC-B. Note that standard molecular weight markers do not accurately reflect MW on Phos-tag gels and were thus not included. (B) Quantitation of densitometry results as shown in (A) and described in the text; results are from six independent determinations from two experiments (C) GC assays of the same six sets of membranes used for (B) under CNP-stimulated conditions. (D-F) Samples from RCS cells virally transduced with WT-GC-B. (E) Quantitation of densitometry results as shown in (D); results from five independent determinations from two blots (F) GC assays of the same five sets of crude membranes used for (E) under CNP-stimulated conditions. **, ***, and **** indicates significance vs. control at p<0.01, 0.001, and 0.0001, respectively, as determined by unpaired, two-tailed student’s t-test.

We also used Phos-tag gel electrophoresis and immunoblotting to investigate the effect of FGF2 on GC-B phosphorylation in RCS cells that were infected with an adenovirus expressing wild type rat GC-B (RCS-GC-B). As with endogenous GC-B, these blots showed a shift in gel migration indicative of dephosphorylation (Fig. 4D). FGF2 decreased the ratio of the intensities of the upper and lower regions by 46% (Fig. 4E). Similar results were obtained using 2 different antibodies made against 2 different epitopes of GC-B (Fig. 4D and Fig. S1), confirming that the observed bands represent GC-B protein.

Guanylyl cyclase assays on the same membrane samples used in Phos-tag experiments indicated that the FGF2-dependent dephosphorylation of GC-B correlated with 74% and 43% reductions in CNP-dependent GC-B activity in membranes from RCS or virally infected RCS cells, respectively (Fig. 4C and F).

Attempts to measure phosphate and protein levels of endogenous GC-B in RCS cells by ProQ Diamond and SYPRO Ruby staining failed because both measurements were below the detection limit of these assays (RCS, Fig. 5A and B). However, these methods were sensitive enough to detected GC-B in the cells infected with an adenovirus overexpressing WT-GC-B (RCS-GC-B). In these cells, FGF2 exposure clearly reduced ProQ Diamond phospho-staining of GC-B without affecting GC-B protein concentrations measured by SYPRO Ruby staining (Fig. 5A and B). Quantitation of the ProQ Diamond/SYPRO Ruby intensity ratio indicated that FGF2 caused a 41% reduction in the phosphate to protein ratio for GC-B (Fig. 5C). GC assays conducted on the same membranes used for the phosphate and protein determinations indicated that FGF2 exposure reduced CNP-dependent, but not detergent-dependent, GC activity by 32%. Since detergent-dependent GC activity did not change, this indicates that the reduction in CNP-dependent activity is not explained by decreases in GC-B protein, and is consistent with inactivation by dephosphorylation.

**Figure 5.**
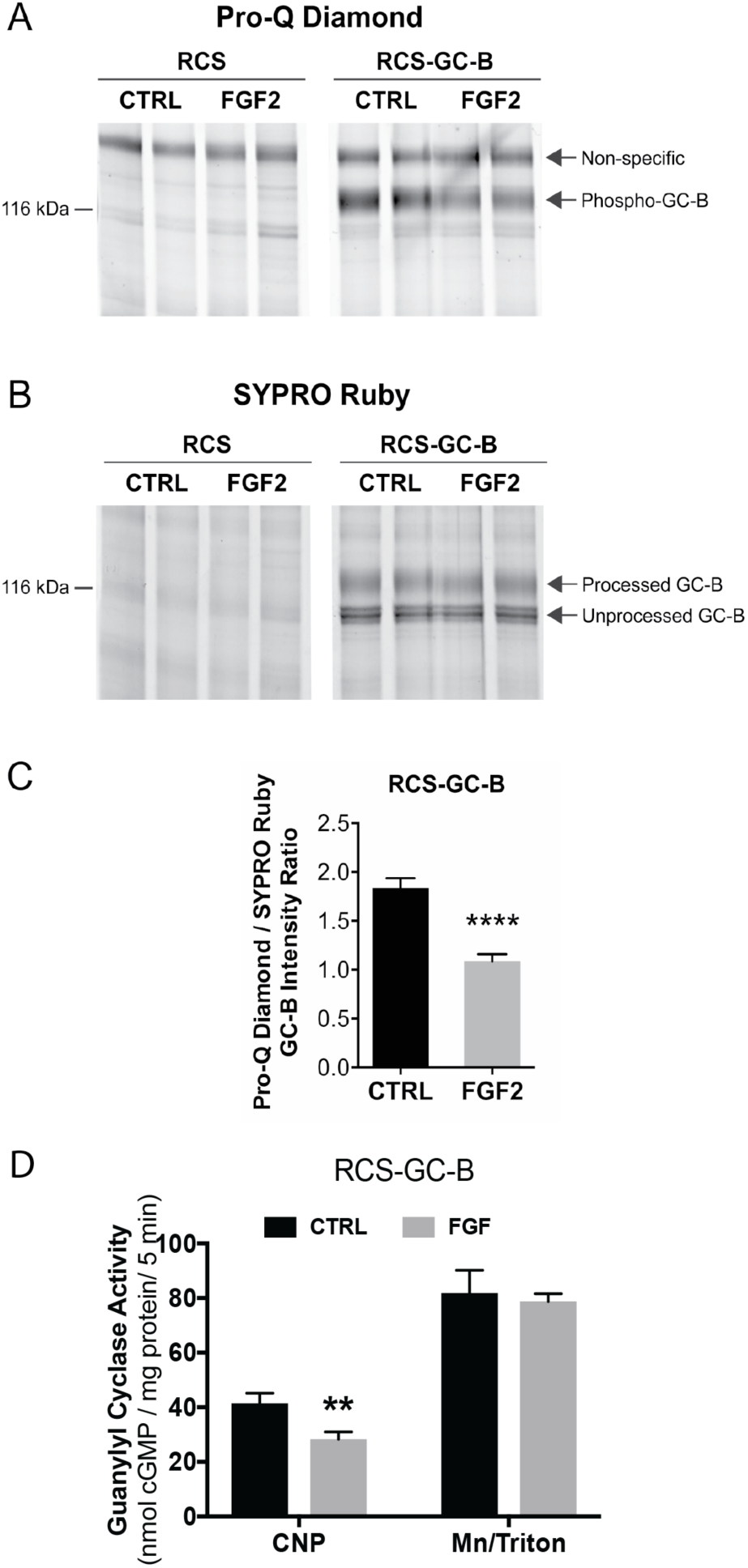
ProQ Diamond staining indicates that FGF exposure causes GC-B dephosphorylation. RCS cells were transduced with an empty virus (RCS) or an adenovirus expressing wild type GC-B (RCS-GC-B). Cells were exposed to 1 μg/ml heparin with or without 100 ng/ml FGF2 for 20 min. Membranes were prepared, immunoprecipitated, and fractionated by SDS-PAGE. The gels were stained with ProQ Diamond followed by SYPRO Ruby. (A and B) Endogenous GC-B in RCS cells was below the detectability limit for ProQ and SYPRO staining (left 4 lanes), but GC-B in virally transduced cells was visible as a band at ∼130 kDa (4 right lanes). (C) Densitometry measurements of the ratio of intensities of the phospho-GC-B (ProQ) and total processed GC-B (SYPRO) bands (results are from 8 determinations and 4 independent experiments). (D) GC assays of the same 8 sets of crude membranes prepared for (C) under CNP-stimulated or detergent-stimulated conditions. ** and **** indicate significance at p<0.01 and 0.0001, respectively, using an unpaired, two-tailed student’s t-test.

To begin to examine which particular phosphatase(s) account for the FGF-induced GC-B dephosphorylation, we tested the effect of the PPP family phosphatase inhibitor cantharidin on FGF-induced dephosphorylation. Phos-tag analysis indicated that 10 μM cantharidin did not inhibit the dephosphorylation (Fig. S2). Cantharidin inhibits PPP1, PPP2, PPP4, PPP5, and PPP6, but not PPP3 or non-PPP family phosphatase [31, 43, 44], so lack of effect of cantharidin on the FGF-induced dephosphorylation suggests that a phosphatase other than PPP1, 2, 4,5, or 6 causes the dephosphorylation. However, we cannot exclude the possibility that the cytoplasmic concentration of cantharidin was insufficient to be effective, or the possibility that cantharidin causes differential phosphorylation of GC-B such that sites that are protected from dephosphorylation cause the dephosphorylation of other sites, resulting in no change in the stoichiometry of GC-B phosphorylation.

### 3.5. FGF-dependent inhibition of GC-B requires receptor dephosphorylation

To test whether dephosphorylation is required for the FGF2 inactivation of GC-B, we created an adenovirus expressing GC-B-7E, a mutant form of GC-B that contains glutamate substitutions for all known and putative phosphorylation sites, which functionally mimics the fully phosphorylated and active enzyme [19]. Unlike the wild type enzyme, GC-B-7E cannot be inactivated by dephosphorylation of the mutated phosphorylation sites [19, 24]. Therefore, we hypothesized that if FGF2 inhibits GC-B through dephosphorylation, then FGF2 should not inhibit GC-B-7E. As predicted, FGF2 failed to affect CNP-dependent GC activity of membranes from GC-B-7E transduced RCS cells (Fig. 6C). ProQ Diamond staining of GC-B-7E revealed a faint band consistent with a previous report showing that proteins with high concentrations of glutamate like lysozyme are slightly stained by ProQ Diamond [42]. Importantly, neither ProQ Diamond nor SYPRO Ruby staining of GC-B-7E was affected by FGF2 exposure (Fig. 6A and B). These data are consistent with a model where dephosphorylation of one or more of the known GC-B phosphorylation sites is required for the FGF2-dependent inactivation of GC-B.

**Figure 6.**
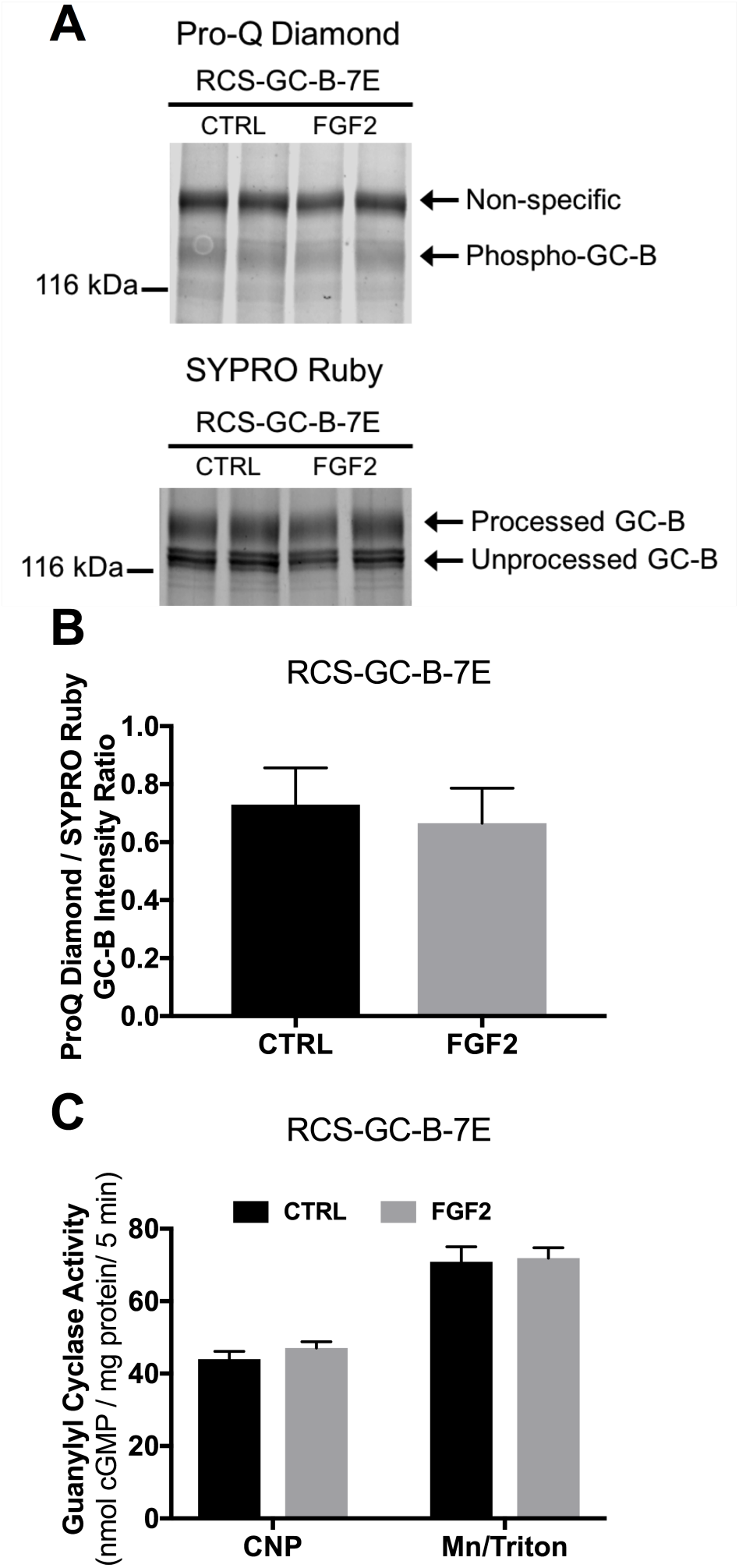
FGF2-dependent GC-B inhibition requires changes in the known GC-B phosphorylation sites. RCS cells were transduced with adenovirus expressing GC-B-7E. Cells were treated with 1 μg/ml heparin with or without 100 ng FGF2 for 20 min. (A) Pro-Q Diamond phosphate staining followed by SYPRO Ruby protein staining of immunoprecipitated GC-B-7E fractionated by SDS-PAGE from one of the sets of membranes assayed for GC activity. The arrow labelled Phospho-GC-B marks the position of phospho-GC-B on gels of wild type GC-B (Fig. 6A); the faint band could represent either staining of glutamate (32) or unidentified GC-B phosphorylation sites that were not mutated to glutamate. The exposure for this image was increased relative to that for Fig. 5A, in order to show the faint band. Correspondingly, the non-specific band is darker in 6A vs 5A. (B) Densitometry measurements of the ratio of intensities of the phospho-GC-B (ProQ) and total processed GC-B (SYPRO) bands. (C) CNP-or detergent-dependent GC activity measurements for the same membranes that were used for the ProQ Diamond and SYPRO Ruby measurements from (B) (results are from 8 determinations that came from 4 independent experiments).

## 4. Discussion

An understanding of phosphorylation-dependent regulation of GC-B has unfolded over the past 20 years. GC-B was first shown to be phosphorylated on serine and threonine residues when purified from stably overexpressing NIH3T3 fibroblasts in 1998 [45]. The receptor was maximally phosphorylated and maximally responsive to CNP stimulation when isolated from serum starved resting cells. In contrast to many G-protein coupled receptors, where prolonged agonist exposure results in receptor phosphorylation and inactivation, the enzymatic activity of GC-B was shown to be highest when maximally phosphorylated. Furthermore, prolonged exposure to CNP resulted in receptor inactivation that was correlated with GC-B dephosphorylation, not phosphorylation [45].

Exposure to hormones or paracrine factors known to antagonize the actions of GC-B by activating G protein coupled receptors also leads to the dephosphorylation [21,25,43] and inactivation [22, 23, 27, 35, 39, 40, 46, 47] of GC-B in other cell types. However, except for a study of luteinizing hormone action on ovarian follicles [25], these previous studies did not couple changes in GC-B phosphorylation to physiological events. The present study links GC-B dephosphorylation and inactivation by a growth factor receptor to mechanisms that regulate bone growth [6,10].

The role of GC-B dephosphorylation in the FGF-dependent inactivation of GC-B has not been reported. Ozasa et al. demonstrated that FGF2 treatment inhibits GC-B in ATDC5 cells and Chrisman and Garbers showed that FGF2 inhibits GC-B in Balb3T3 cells, but neither group determined the mechanism of the inactivation [26, 27]. Here, we report that FGF2 activation of FGFR3 in RCS cells results in a rapid, potent, and reversible inhibition of CNP-dependent GC-B activity. The reversibility and speed of FGF2-dependent inhibition indicate that GC-B degradation is not required for the inhibition and are consistent with a rapidly reversible process like dephosphorylation. The fact that SYPRO Ruby staining of GC-B did not change in response to FGF2 exposure also indicates that the loss of activity is independent of changes in GC-B protein. The potency data suggested that the FGF2-dependent inhibition is physiologically relevant and is a potential regulatory mechanism for GC-B *in vivo*. Importantly, GC-B dephosphorylation was measured using two different phosphorylation detection methods. Furthermore, the inability of FGF2 to inhibit a dephosphorylation-resistant form of GC-B (GC-B-7E) provides a critical test of the “inactivation by dephosphorylation” hypothesis and is direct evidence demonstrating that prevention of FGF-dependent dephosphorylation of known phosphorylation sites prevents GC-B inactivation.

In conclusion, these data in combination with previously described reports of GC-B being inactivated by dephosphorylation in response to other signaling molecules, strongly suggests that distinct hormonal systems inactivate GC-B by a general dephosphorylation mechanism. Identification of the modified GC-B phosphorylation sites will be an important next step in defining the molecular basis for how hormones and growth factors inhibit GC-B. RCS cells overexpressing GC-B are a promising system for addressing this issue, since physiological responsiveness to FGF is retained under conditions where GC-B protein levels are elevated about ten-fold over endogenous levels. Also, identifying the specific downstream signaling pathways required for the FGF2 inhibition of GC-B will be important to understand how FGF2 inhibits GC-B. Additionally, it will be interesting to determine whether long bone growth and/or long bone composition is changed in GC-B^7E/7E^ compared to GC-B^WT/WT^ mice or whether mice expressing dephosphorylated forms of GC-B are dwarfed.

Finally, we suggest that understanding the cross-talk between the FGFR3 and GC-B signaling cascades is of increasing medical importance because a protease resistant analog of CNP is in clinical trials as a therapy for increasing longitudinal growth in children with dwarfism [48, 49]. However, a recent report showing higher circulating levels of CNP in children with achondroplasia suggests tissue resistance to CNP [50], which may result from increased dephosphorylation and desensitization of GC-B. In either instance, understanding the contribution of GC-B dephosphorylation and desensitization to achondroplasia is necessary to ensure drugs that specifically target GC-B will be effective therapeutics for achondroplasia.

## Acknowledgments

We thank Kari Jacobsen for expert preparation of the GC-B-WT and GC-B-7E adenoviruses. This work was supported by National Institutes of Health Grants, R01GM098309 to LRP, R37HD014939 to LAJ, T32DK007203 to JWR, by a German Research Foundation (DFG) grant (FOR 2060, HSCHM 2371/1) to HS and by the Fund for Science and the Hormone Receptor Fund.

---

**Fig. S1.**
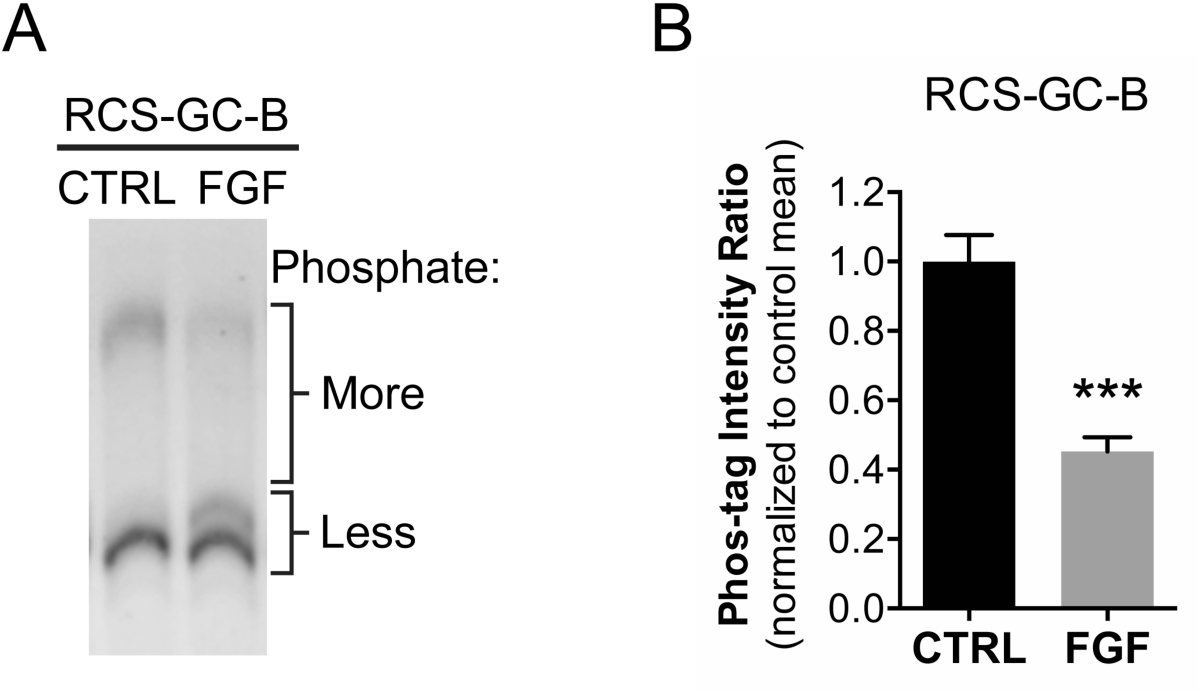
Control for antibody specificity using 2 different primary antibodies for probing the same Western blot. (A) The same blot shown in Fig. 4D was probed with antiserum against the C-terminus of GC-B (6328) instead of the antibody against the extracellular domain of GC-B that was used for the image in Fig. 4D. The blot was stripped between probing with the 2 different antibodies by immersing in buffer containing 62.5 mM Tris-HCl pH 6.8, 100 mM 2-mercaptoethanol, and 2% SDS for 30 min at 50°C, followed by extensive washing and blocking as described in the text. (B) Quantitation of the Phos-tag intensity ratio (upper region/lower bands) in control and FGF-treated cells, normalized to the mean of the control lanes. Results from 5 independent experiments on 2 blots. *** indicates significance vs. control at p<0.001 using an unpaired two-tailed student’s t-test.

**Figure S2.**
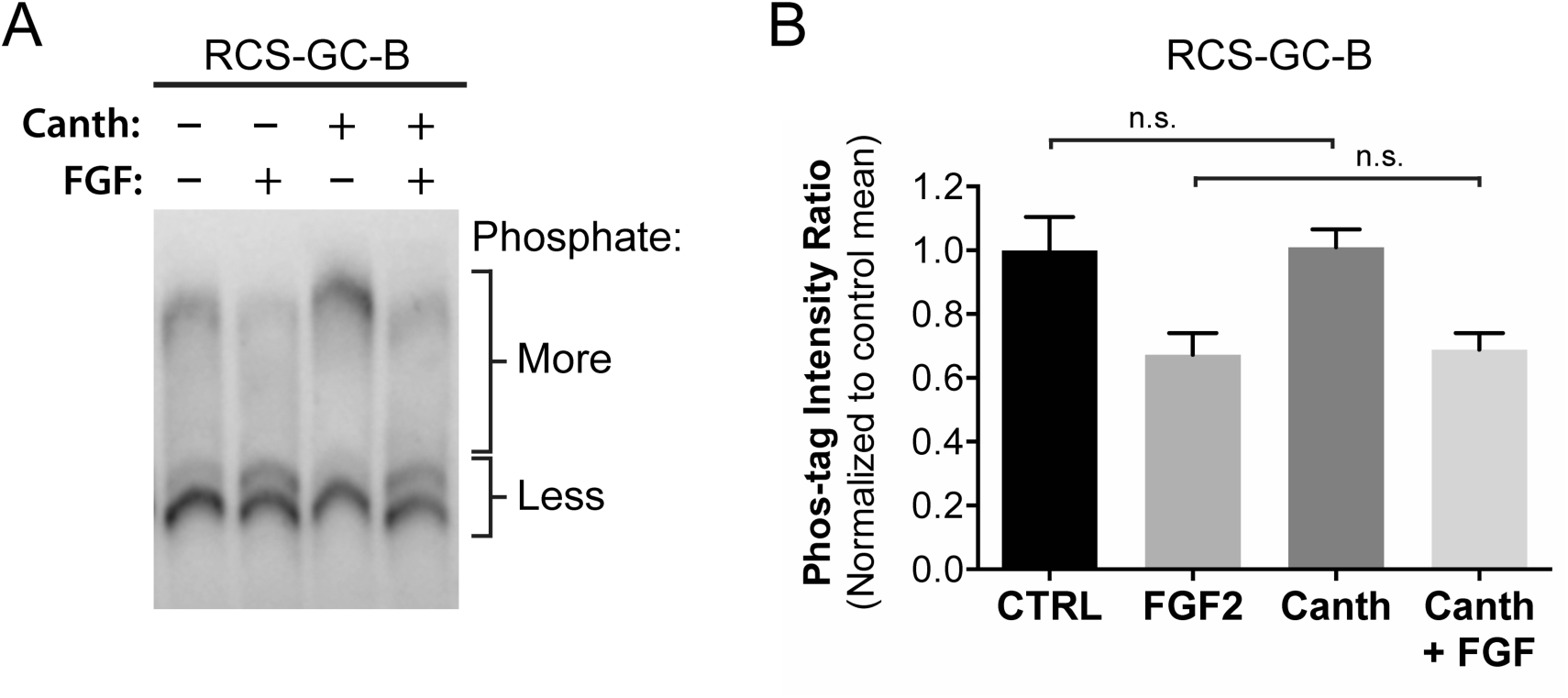
Lack of effect of cantharidin on FGF-induced dephosphorylation of GC-B. (A) RCS-GC-B cells were pretreated for 1 hour with or without 10 μM cantharidin, then for 20 min with 1 μg/ml heparin with or without 100 ng/ml FGF2. Crude membranes were prepared, and GC-B was immunoprecipitated using an antibody against the C-terminus of GC-B [35]. The immunocomplexes were separated by Phos-tag gel electrophoresis, then proteins from the gel were transferred to a membrane for immunoblotting. The blot was probed with an antibody against the extracellular domain of GC-B. Lanes 1 and 2 are the same as those shown in Fig. 4A. (B) Quantitation of the Phos-tag intensity ratio (upper region/lower bands) in control and treated cells, normalized to the mean of the control lanes. Results from 3 independent experiments, analyzed on the same blot. Cantharidin had no effect on GC-B phosphorylation either without or with FGF treatment; n.s. indicates p>0.05 using unpaired two-tailed student’s t-tests.

